# A Method for Predicting Enzyme Substrate Specificity Residues Using Homologous Sequence Information

**DOI:** 10.1101/2025.05.25.656053

**Authors:** Seiya Mori, Teppei Niide, Yoshihiro Toya, Hiroshi Shimizu

## Abstract

Identifying amino acid residues that are critical for the catalytic function of enzymes is essential for elucidating reaction mechanisms, facilitating drug discovery, and advancing protein engineering. However, experimentally and computationally distinguishing residues that maintain structural integrity from those directly involved in enzymatic function remains a major challenge. In this study, we developed a methodology to identify amino acid residues that influence substrate specificity in enzymes with homologous structures. We framed the sequence comparison as a classification problem, treating each residue as a feature, thereby enabling the rapid and objective identification of key residues responsible for functional differences. To validate the proposed method, we applied it to three enzyme pairs— trypsin/chymotrypsin, adenylyl cyclase/guanylyl cyclase, and lactate dehydrogenase (LDH)/malate dehydrogenase (MDH). The results confirmed the accurate prediction of previously identified specificity-determining residues. Furthermore, we conducted experiments on the LDH/MDH pair and successfully introduced mutations into key residues to alter substrate specificity, enabling LDH to utilize oxaloacetate while maintaining its expression levels. These findings demonstrate the potential of this method for efficiently identifying residues that govern substrate specificity. We have further developed this approach into a practical tool, the EZSCAN: Enzyme Substrate-specificity and Conservation Analysis Navigator (https://ezscan.pe-tools.com/), which enables rapid identification of amino acid residues critical for enzyme function.

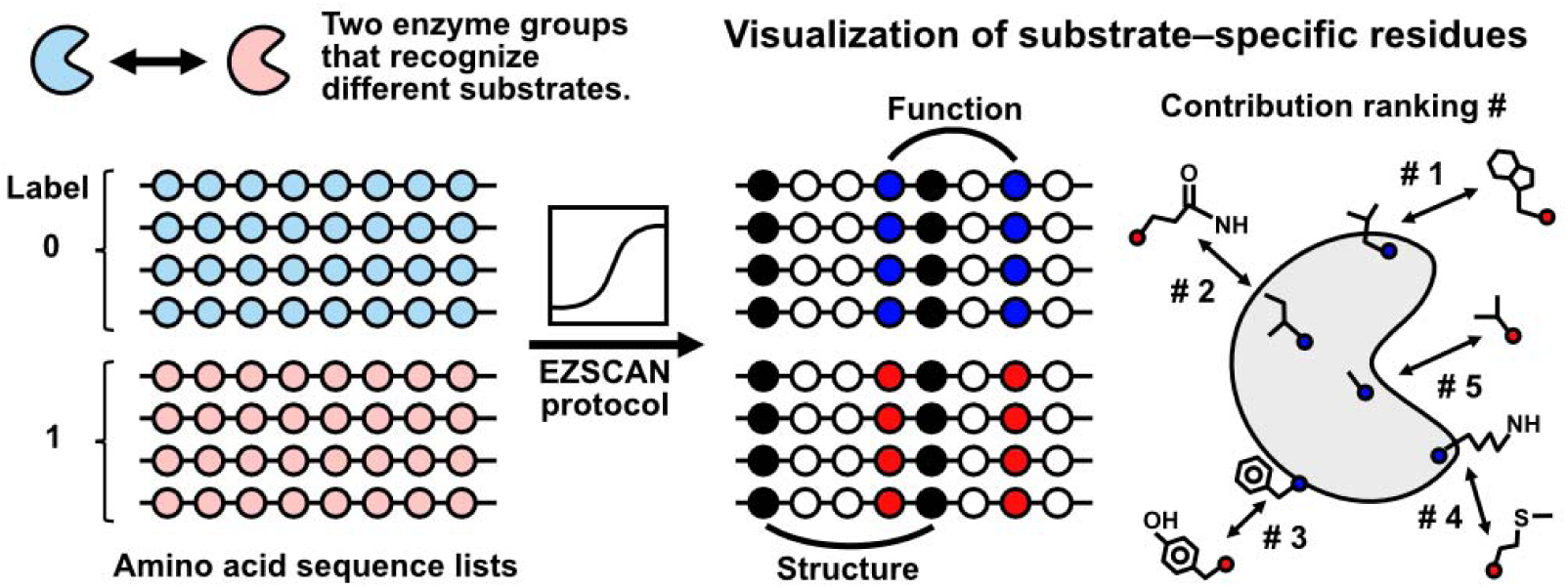

## INTRODUCTION

Enzymes are biological catalysts that drive chemical reactions with high precision and efficiency. Their catalytic function originates from coordinated conformational changes and electron transfer mediated by amino acid side chains^1-4^. Identifying the amino acid residues essential for catalysis and substrate specificity is critical for understanding structure–function relationships. This knowledge provides practical insights into how mutations contribute to disease pathogenesis and how targeted mutagenesis can be used to redesign enzyme specificity for biotechnological applications, such as metabolic engineering and cell manipulation^5, 6^.

Advances in DNA synthesis and sequencing technologies have paved the way for in-depth studies of mutation–function relationships, such as deep mutational scanning and multiplexed variant assays^7-9^. These methods systematically mutate each amino acid residue into all 20 possible variants, generating extensive libraries. Using NGS, researchers can identify residues critical for enzyme function by subjecting these populations to functional selection and tracking changes in abundance. Although large-scale mutation analysis offers functional mutation profiles, distinguishing functionally critical residues from those conserved due to structural constraints remains a challenge, as does identifying synergistic mutations^10^. Moreover, as nearly half of the loss-of-function mutations result from decreased protein abundance^11^, new approaches are necessary to pinpoint residues directly influencing enzyme function.

In contrast to experimental methods, computational techniques offer rapid, cost-effective, and scalable alternatives for identifying functionally important residues. Several computational approaches have been developed to predict these residues based on their amino acid sequences. Among these, conservation analysis—which identifies residues that are highly conserved across proteins—has emerged as a powerful tool for identifying functionally critical residues^12-15^. Although this method has provided valuable insights into protein–protein interactions, structural stability, and ligand recognition, it requires refinement to distinguish between residues essential for function and those conserved due to structural constraints. As both functional and structural constraints shape protein evolution, the challenge is to identify residues that are crucial for protein function without being confounded by structural conservation.

Recent advances in molecular biology and machine learning have shed light on how specific amino acid residues determine cofactor specificity. Using supervised learning on amino acid sequence datasets, we previously identified key residues that distinguish between NAD(H)- and NADP(H)-dependent malic enzymes^16^. Despite clear differences in cofactor preferences, these enzymes retain a highly conserved overall structure across species. Guided by machine learning-based residue rankings, we introduced mutations that not only preserved soluble expression but also completely switched the enzyme’s cofactor specificity from NADP to NAD. Notably, these substitutions were well tolerated, underscoring the functional relevance of the identified sites and enabling the separation of structural and functional constraints, which is difficult to achieve through conventional conservation analysis. These findings pointed to a broader principle: functionally critical residues underlying substrate specificity can be identified by contrasting enzymes that are structurally conserved yet functionally distinct.

In this study, we present a computational framework to uncover the molecular basis of enzyme substrate specificity. By analyzing the sequence datasets of homologous enzymes using supervised machine learning, we identified key amino acid residues that govern substrate recognition. Focusing on three well-studied enzyme pairs—trypsin/chymotrypsin, adenylyl/guanylyl cyclase (AC/GC), and lactate/malate dehydrogenase (LDH/MDH)—we recovered known specificity-conferring residues and revealed previously unreported sites critical for function. Experimental validation of the LDH/MDH pair confirmed that the newly identified residues contributed directly to differences in substrate preference. Although protein sequences and functions have diversified through evolution, their three-dimensional structures are often highly conserved^17, 18^. Comparative analyses of these protein families can provide key insights into the evolution of functions while maintaining structural constraints. Our approach not only distinguishes functionally relevant positions in structurally similar enzymes but also provides a generalizable strategy for dissecting enzyme specificity. To support its broad adoption, we developed EZSCAN, a web tool that enables researchers to explore substrate recognition features across diverse enzyme families.

## RESULTS

### Prediction of substrate-specific residues in enzymes using EZSCAN

The EZSCAN protocol represents an advancement in understanding enzyme functionality through a machine-learned binary classification algorithm tailored to extract critical amino acid residues linked to enzyme cofactor specificity. In this study, we extended this approach to identify amino acid residues vital for substrate specificity by leveraging two distinct sets of amino acid sequence data (Figure 1). Initially, we obtained amino acid sequences of two sets of enzymes with homologous structures from a comprehensive database. These sequences were aligned using multiple sequence alignment, converted into one-hot vectors, and subsequently analyzed using a logistic regression model. Given the structural homology between the enzyme sets, our classification of residues associated with enzymatic function is expected to yield meaningful results. The key explanatory variable in the trained model was the amino acid type at each position, and the range between the maximum and minimum partial regression coefficients served as an important evaluative metric.

**Figure 1.**
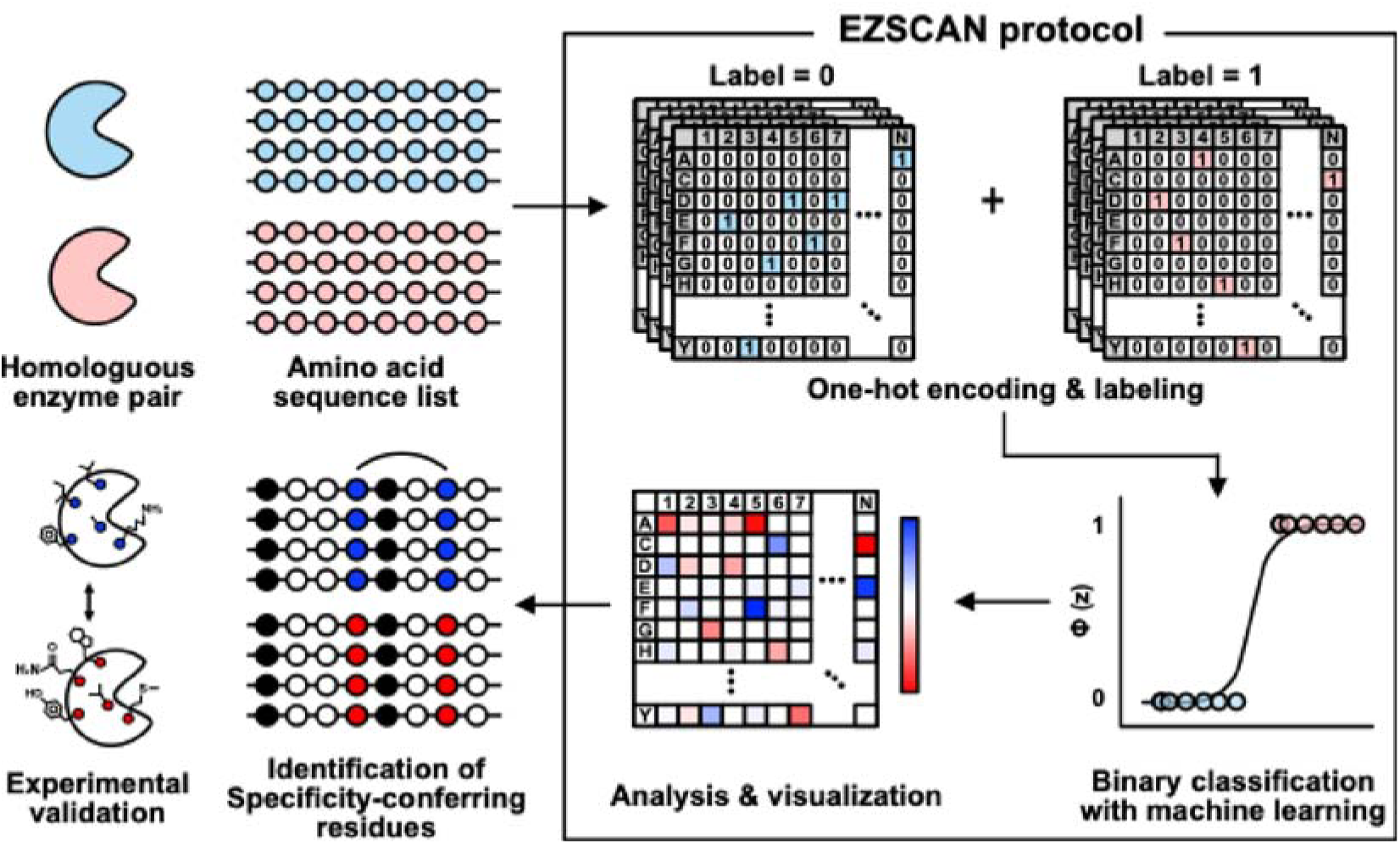
Schematic illustration of estimation process for specificity-conferring residues using the EZSCAN protocol. The amino acid sequences of two structurally homologous enzyme groups are used as input data, and a logistic regression model is trained to extract specificity-conferring residues that distinguish the two enzyme groups. The extracted specificity-conferring residues can be ranked by their contribution and used for experimental evaluation.

As a practical demonstration, we applied this method to three enzyme pairs— trypsin/chymotrypsin, AC/GC, and LDH/MDH—to identify residues critical for enzymatic function. Despite their differences in substrate preference, these enzyme pairs share homologous structures (Figure 2). The average root mean square deviation (RMSD) and template modeling score (TM-score) values for each enzyme pair further confirmed their high degree of structural similarity. Details of RMSD and TM scores across all crystal structures are provided in the Supplementary Information (Supplementary Tables S1–S6). TM-scores range from 0 to 1, with values above 0.5 indicative of structural homology^19^. In the following sections, we present results identifying amino acid residues that are pivotal for substrate specificity across these different enzymes.

**Figure 2.**
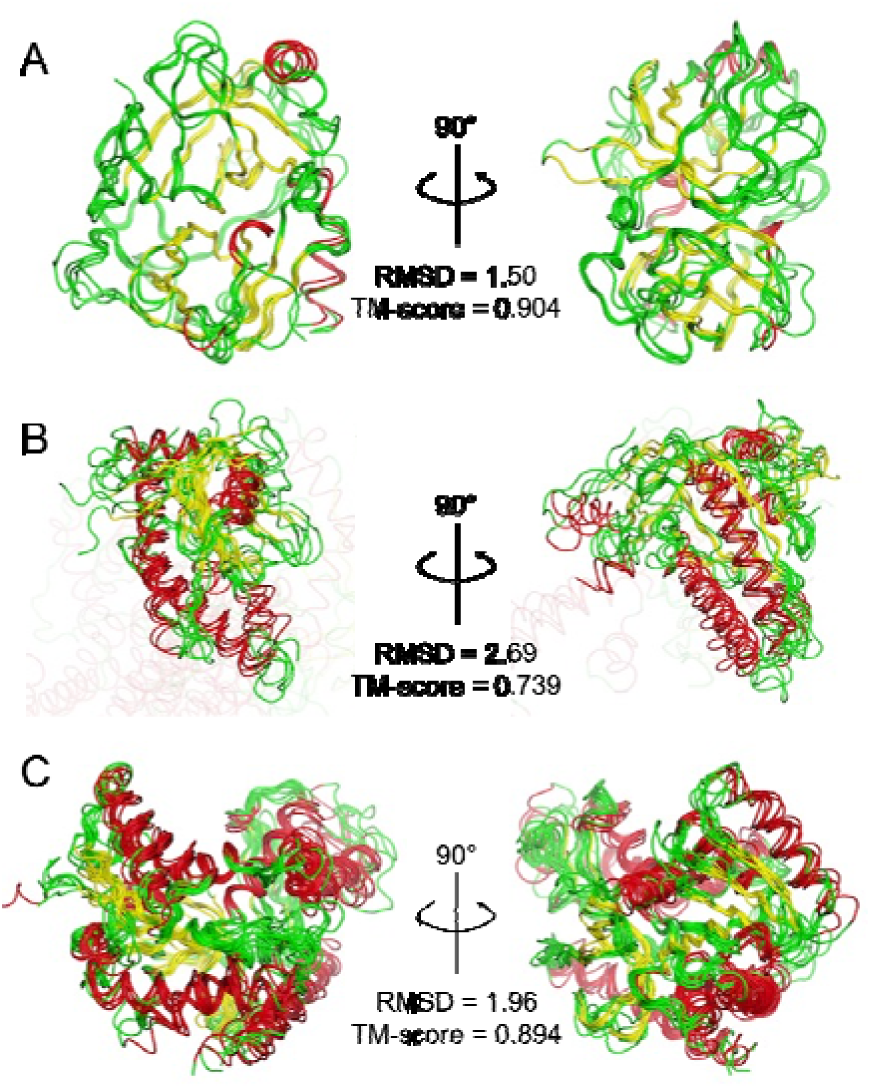
Superimposed images. (A) Trypsin (PDB ID 6T5W, 1ANE, and 1OS8) and chymotrypsin (PDB ID 1KDQ, 1ACB, and 1EQ9). (B) AC (PDB ID 1AB8, 6R3Q, and 7YZI) and GC (PDB ID 2WZ1, 3ET6, and 6PAS). (C) LDH (PDB ID 1LDG, 1LDN, 3VPH, 4AJ2, and 6J9T) and MDH (PDB ID 1B8P, 1HLP, 2PWZ, 4CL3, and 5UJK). Red, yellow, and green regions represent α-helix, β-sheet, and random coil structures, respectively. Average values of metrics from the superimposed structures are shown. All scores compared between each structure are shown in Supplementary Tables S1–S6.

#### Trypsin/chymotrypsin

Trypsin and chymotrypsin, both serine proteases, exhibited significant structural homology (Figure 2A). Trypsin is known for cleaveing the C-terminal side of Arg and Lys at the P1 position of substrate peptides, whereas chymotrypsin targets Phe, Tyr and Trp at the same position^20^. This distinction in substrate specificity is primarily attributed to the pivotal role of residue S195 (in chymotrypsin numbering), which defines the S1 pocket adjacent to the active site^21^. Notably, the negative charge generated by the combination of D189, G216, and G226 (chymotrypsin numbering) significantly influences trypsin substrate specificity. In contrast, S189, G216, and G226 (chymotrypsin numbering) are critical for chymotrypsin specificity ^22^. Interestingly, swapping D189 and S189 alone is insufficient to shift specificity from trypsin-like to chymotrypsin-like or vice versa ^23, 24^. Additionally, Y172, although not directly involved in substrate interaction, is essential when transitioning substrate specificity between two enzymes ^25^. To explore the predictive potential of EZSCAN for identifying specificity-conferring residues, we applied the protocol to trypsin and chymotrypsin.

Amino acid sequence data for these enzymes were obtained from the Kyoto Encyclopedia of Genes and Genomes (KEGG), focusing on sequences between 240–270 residues in length. A total of 793 trypsin sequences and 652 chymotrypsin sequences were used as input for EZSCAN analysis (Supplementary Figure S1A). We used trypsin and chymotrypsin structures from *Rattus norvegicus* as templates to display specificity-conferring residues. The model predicted Tyr (trypsin) and Trp (chymotrypsin) at residue 172 as the top specificity-conferring residues (Figure 3A, Table 1). Notably, Asp (trypsin) and Ser (chymotrypsin) at residue 189 were ranked fourth, consistent with prior studies on substrate specificity.

**Figure 3.**
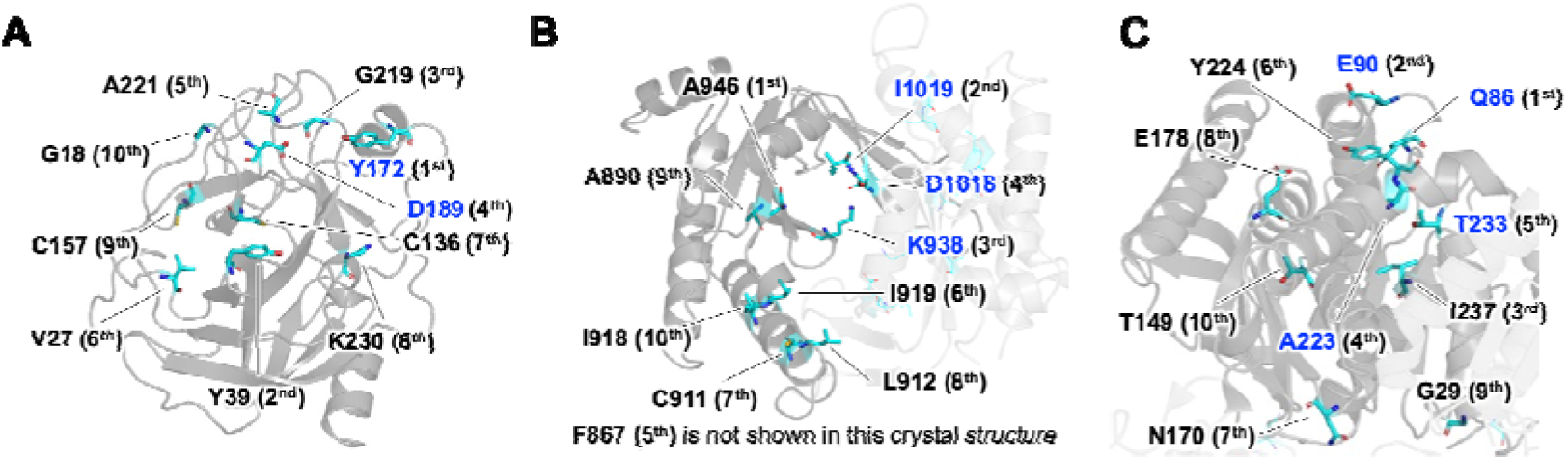
Mapping of specificity-conferring residues estimated by EZSCAN on the crystal structure. (A) Trypsin derived from *R. norvegicus* (PDB: 1ANE), (B) AC derived from *R. norvegicus* (PDB: 1AB8), (C) LDH derived from *G. stearothermophilus* (PDB: 1LDN). Cyan sticks represent the top 10 ranked amino acid residues estimated by the EZSCAN protocol. Residues previously reported to be involved in substrate specificity are shown in blue; all others are shown in black. The numbers in parentheses indicate their ranking.

**Table 1.**
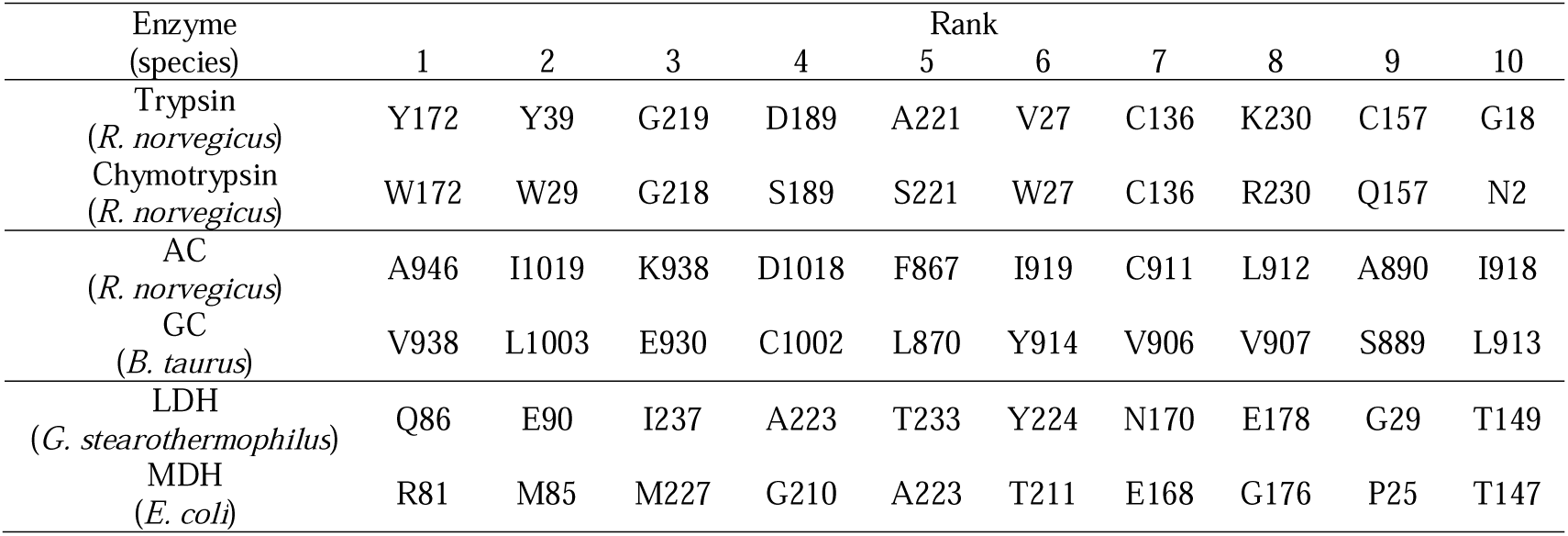
Top 10 ranked amino acid residues associated with substrate specificity for trypsin/chymotrypsin, AC/GC, and LDH/MDH as predicted by EZSCAN. The predicted residue positions correspond to the amino acid positions in each template enzyme. For trypsin/chymotrypsin, the residue numbering follows the chymotrypsin numbering scheme.

The second ranked residue, Tyr (trypsin) and Trp (chymotrypsin) at residue 39, is located distally from the active site but is conserved in mesotrypsin, which is known to interact with protease inhibitors ^26^. This suggests that the second ranked residue may also influence substrate recognition. Additionally, the third ranked residue at position 219 and the fifth ranked residue at position 221 form a loop near the substrate pocket, indicating their likely contribution to the substrate recognition process.

#### AC/GC

AC and GC are crucial enzymes in signal transduction, responsible for converting ATP and GTP into cAMP and cGMP, respectively. ACs are classified into five categories based on their structural characteristics^27, 28^. Notably, Class III ACs exhibit a close phylogenetic relationship with GCs, and their catalytic domains demonstrate significant structural homology (Figure 2B). The catalytic domain of mammalian Class III AC exists as a heterodimer composed of C1 and C2 domains, whereas the bacterial and protozoan counterparts are homodimeric^29, 30^. These enzymes exhibit strict selectivity for ATP or GTP, with specific amino acid residues at the dimer interface playing a pivotal role in determining substrate specificity^31-33^. For instance, it has been shown that GC from *Bos taurus* can be engineered to function similarly to AC through just two amino acid substitutions at the active site—E930K and C1002D^34^. In contrast, similar modifications at corresponding sites in AC from *R. norvegicus* did not produce a substantial change in substrate specificity.

Residue I1019 has been implicated in the formation of hydrogen bonds with the N-6 amino group of ATP and is considered essential for this process^35^. We aimed to evaluate whether such specificity-conferring residues in AC and GC could be predicted using the EZSCAN software.

Amino acid sequence data for AC and GC were obtained from the KEGG database. We analyzed 319 ACs and 572 GCs, with sequence lengths ranging from 1,090 to 1,130 amino acids (Supplementary Figure S1B). In this analysis, AC from *R. norvegicus* and GC from *B. taurus* served as template structures for identifying specificity-conferring residues. EZSCAN identified E930 and C1002 as the third and fourth highest-ranked residues for AC from *R. norvegicus*, consistent with previous findings demonstrating their role in shifting substrate specificity (Figure 3B, Table 1). Another key residue, I1019, ranked second and is known to directly interact with the substrate.

The top-ranked mutation in GC has been implicated in central areolar choroidal dystrophy, a genetic eye disease ^36^, indicating its potential significance in cyclase function. Interestingly, although our analysis encompassed the full-length sequences of both AC and GC, the top-ranked residues predominantly resided in the C2 domain. This finding aligns with prior studies^34, 35, 37, 38^ and suggests that the presence of non-homologous transmembrane domains does not interfere with the identification of residues essential for substrate specificity.

#### LDH/MDH

LDH and MDH are essential redox enzymes characterized by structural homology and a reliance on NAD(H) as a cofactor (Figure 2C). LDH catalyzes the interconversion between lactate and pyruvate, while MDH is involved in the conversion of malate and oxaloacetate^39, 40^. The differences in substrate specificity between these enzymes are primarily due to variations in amino acid residues within their substrate-binding sites. Notably, LDH from *Geobacillus stearothermophilus* can acquire MDH activity through a single Q86R mutation ^41^. Conversely, introducing the reverse mutation into MDH from *Escherichia coli* does not result in a comparable enhancement in LDH activity^42, 43^. Furthermore, one study reported that introducing five specific mutations—I12V, R81Q, M85E, G210A, and V214I—dramatically increased the *k_cat_*/K_M_ value of *E. coli* MDH from 0.14 to 3,500 MlJ¹slJ¹ ^44^. These findings raise an important question: can EZSCAN accurately identify the residues responsible for such differences in substrate specificity between LDH and MDH?

We sourced amino acid sequences for LDH and MDH from the UniProtKB database and analyzed 228 LDH and 397 MDH sequences. The sequence lengths ranged from 300 to 340 amino acids (Supplementary Figure S1C). Using LDH from *G. stearothermophilus* and MDH from *E. coli* as template structures, we identified key specificity-conferring residues. Residues Q81, M85, and G210, previously reported to play crucial roles in substrate specificity, were ranked first, second, and fourth, respectively (Figure 3C, Table 1). The fifth-ranked mutation, from Thr to Gly, was shown to reduce the enzymatic activity of *G. stearothermophilus* LDH by more than 1,000-fold, highlighting its importance in substrate recognition^41^. Interestingly, the third-ranked residue, I237, has not been previously reported but is located at the base of the substrate pocket, suggesting a potential role in influencing substrate specificity.

We applied the EZSCAN protocol to three enzyme pairs based on prior experimental findings. This approach successfully identified amino acid residues known to determine substrate specificity and ranked them prominently. Additionally, EZSCAN revealed new candidate residues that have not been examined previously, offering valuable targets for future experimental validation. This method also provides a comprehensive view by quantifying the contribution of all 20 amino acid variants at each residue position, enabling a clearer understanding of the mechanisms underlying enzyme specificity (Supplementary Figures S2–S4). These findings support the conclusion that machine learning analysis of homologous enzyme sequences can effectively uncover substrate specificity-conferring residues.

### Selection of LDH for Experimental Validation

We experimentally validated the amino acid residues associated with substrate specificity, as predicted by the EZSCAN protocol. LDH catalyzes the interconversion between pyruvate and lactate, whereas MDH catalyzes the conversion of oxaloacetate to malate (Figure 4A). To evaluate the accuracy of EZSCAN predictions, we assessed whether the specificity of template LDHs for pyruvate could be decreased and their specificity for oxaloacetate increased.

**Figure 4.**
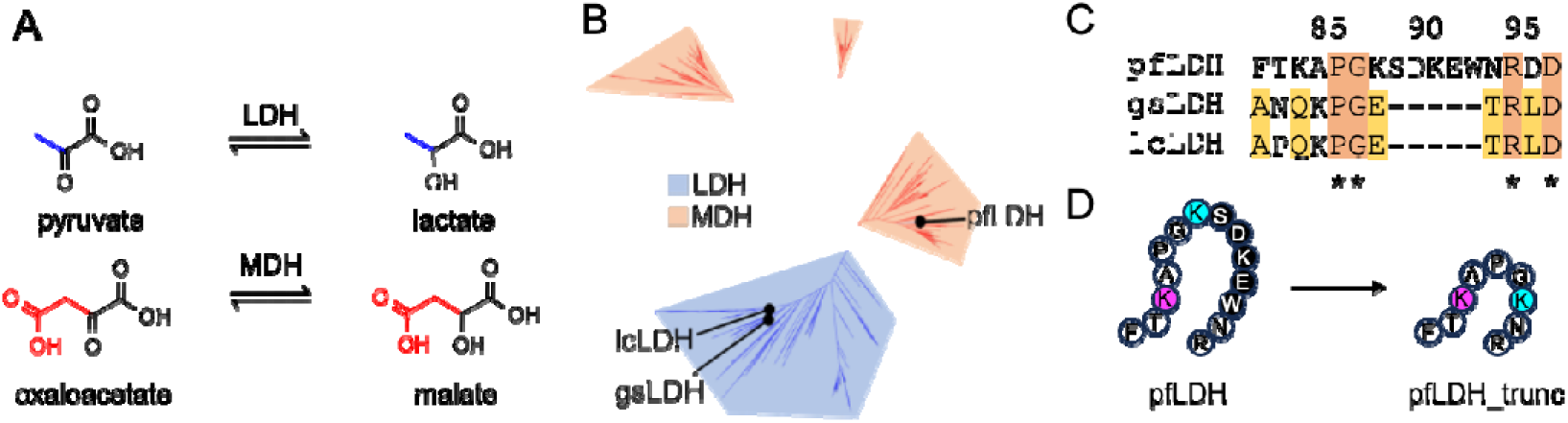
(A) Reactions catalyzed by LDH and MDH. (B) Phylogenetic tree constructed from the LDH/MDH dataset used for model training. Blue and orange branches represent LDH and MDH, respectively. (C) Aligned amino acid sequences of pfLDH, gsLDH, and lcLDH. Dashes indicate alignment gaps. An insertion of SDKEW is observed at positions 89–93 in pfLDH. Residue numbering corresponds to pfLDH. (D) Design of pfLDH_trunc, in which residues 89–93 (SDKEW) of pfLDH are deleted. Black indicates the truncated region; magenta and cyan highlight residues ranked first and second in importance, respectively.

Four LDHs from three species were used as template enzymes, and the EZSCAN-predicted specificity-conferring residues in these LDHs were replaced with MDH-like residues. The LDHs selected for mutagenesis were LDH from *G. stearothermophilus* (gsLDH; UniProt ID: P00344), LDH from *Lactobacillus casei* (lcLDH; UniProt ID: P00343), and LDH from *Plasmodium falciparum* (pfLDH; UniProt ID: Q27743). gsLDH has previously been reported to acquire MDH activity via a single Q86R mutation ^41^. lcLDH was selected because it has a lower optimal pH (4.8) than other LDHs ^45^, which could lead to distinct mutational effects compared with gsLDH.

pfLDH is a unique LDH found within the MDH clade rather than the LDH clade (Figure 4B), suggesting it may have arisen by convergent evolution from an MDH ancestor ^46^. Its sequence is therefore expected to resemble that of MDH, making it a suitable candidate for examining differences between enzyme templates. Alignment of pfLDH with gsLDH and lcLDH revealed a five-amino acid insertion from S89 to W93 in pfLDH (Figure 4C). These inserted residues are located in the same loop where EZSCAN ranked residues first and second in importance, suggesting that the insertion likely alters the shape of the substrate pocket (Supplementary Figure S5).

In addition to wild-type pfLDH, we designed a truncated variant, pfLDH_trunc, in which the five-residue insertion was removed (Figure 4D). Structural prediction indicated that pfLDH_trunc is homologous to gsLDH and lcLDH (Supplementary Figure S6). Using these four types of LDHs as templates, we evaluated whether the EZSCAN-predicted specificity-conferring residues, when replaced, would alter substrate specificity.

### Experimental Validation of Substrate Specificity Conversion

The investigation of four types of LDH involved substituting residues identified by the EZSCAN protocol with MDH-like amino acid residues to assess their impact on substrate specificity. These substitutions were introduced sequentially, one residue at a time, starting with the highest-ranked candidate. Because pfLDH retained its third-ranked Pro residue, pfLDH3 incorporated the fourth-ranked amino acid, pfLDH4 added the fifth-ranked residue, and pfLDH5 introduced the sixth-ranked residue.

Each gene encoding the wild-type LDH and its corresponding mutants was cloned into a pET28a expression vector. Expression was carried out in *E. coli* BL21(DE3), and purification was performed using a Ni-NTA column. All LDH variants, including the mutants, were obtained in the soluble fraction. Expression levels ranged from 40.0 to 167.2% relative to the wild-type pfLDH, with no significant decrease in expression observed (Supplementary Figure S6).

To analyze substrate specificity, the purified enzymes were tested using pyruvate and oxaloacetate, and initial reaction velocities were determined by monitoring NADH oxidation at 340 nm. Kinetic parameters were derived from these initial velocities via nonlinear fitting, as shown in Figure 5. Notably, the Q86R mutation in gsLDH (gsLDH1) drastically reduced its LDH activity, decreasing its *k_cat_*/K_M_ value to 1/438 of the wild-type level. At the same time, it enabled MDH activity that was undetectable in the wild-type enzyme. gsLDH1 exhibited a *k_cat_*/K_M_ for MDH of 366.1 mMlJ¹ slJ¹, surpassing the wild-type LDH activity. This switch in substrate specificity due to the Q86R mutation is consistent with previous findings ^41^. Further mutations in gsLDH3 caused LDH activity to fall below detection limits, yielding a purely MDH-like enzyme. gsLDH4 showed a 4.4-fold increase in MDH activity compared to gsLDH3, while gsLDH5 lost MDH activity entirely.

**Figure 5.**
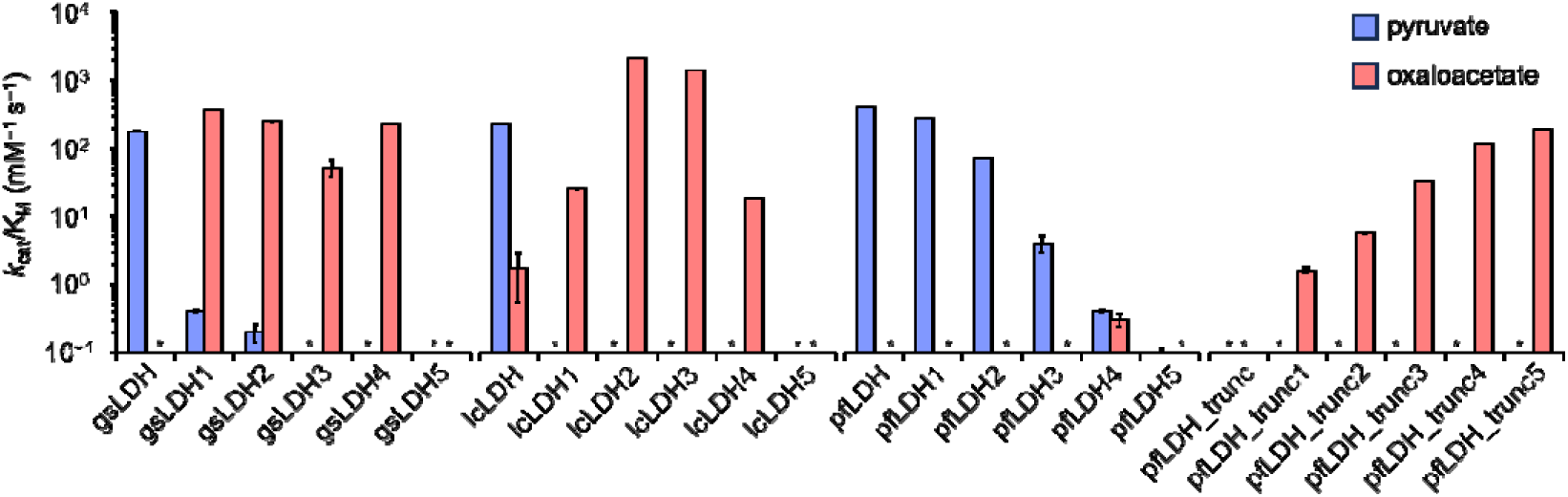
Enzyme activity (*k_cat_*/K_M_) of each LDH variant before and after mutation, based on rankings from the EZSCAN protocol. Blue bars represent enzyme activity using pyruvate as the substrate, and red bars represent activity using oxaloacetate. Data were collected in triplicate. Asterisks indicate cases in which *k_cat_*/K_M_ values could not be determined. Michaelis–Menten plots and fitted curves are provided in Supplementary Figure S7. The *k_cat_* and K_M_ values for each LDH are listed in Supplementary Table S7.

In contrast, lcLDH lost LDH activity with just a single mutation (Q88R), while showing a 15-fold increase in MDH activity, which had previously been nearly undetectable. The lcLDH2 variant, incorporating the additional second-ranked mutation (E92M), displayed an 80-fold increase in MDH activity—a 1,191-fold increase over the wild-type. However, as more mutations were added, MDH activity gradually declined, and lcLDH5 ultimately lost all MDH activity.

For pfLDH, a consistent decline in LDH activity was observed with each successive mutation, ranked according to EZSCAN. pfLDH5 retained only 1/3987 of the wild-type LDH activity. Although pfLDH4 exhibited slight MDH activity, the conversion to MDH-like specificity was not as pronounced as in gsLDH or lcLDH. These results suggest that the SDKEW insertion sequence, spanning residues 159–163 in pfLDH, hinders the acquisition of MDH activity while preserving LDH function.

We further evaluated pfLDH_trunc, in which the SDKEW sequence was removed. This truncation led to a complete loss of LDH activity, consistent with previous findings by Douglas et al. ^46, 47^. This outcome underscores the importance of the SDKEW loop structure in maintaining LDH activity. Interestingly, pfLDH_trunc gained progressively stronger MDH activity with the introduction of additional mutations. Among these, pfLDH_trunc5 exhibited the highest MDH activity. This result suggests that converting pfLDH to MDH functionality—likely governed by a distinct reaction mechanism— requires both structural adjustments to the backbone near functional residues and targeted substitutions.

These findings validate that EZSCAN-predicted residues directly influence substrate specificity. All introduced mutations led to reductions in LDH activity while promoting MDH activity, without significantly affecting expression levels. This confirms that the amino acid residues identified by EZSCAN are indeed specificity-conferring residues.

## DISCUSSION

Directed evolution is a widely employed and powerful strategy in protein engineering. However, the vast sequence space of proteins makes it challenging to efficiently identify variants with the desired functions efficiently ^48^ In typical directed evolution, random mutations or DNA shuffling are introduced into natural proteins, followed by high-throughput screening under selective pressure to enhance protein properties such as activity or stability. Although the number of substitutions required to achieve a functional shift varies depending on the direction of engineering, a few to a dozen amino acid changes are often sufficient for functional enhancement or adaptation. Importantly, protein function and stability often exhibit a trade-off relationship ^49, 50^, highlighting the need for strategies that improve function without compromising structural integrity. A key to the successful design for substrate specificity conversion is to distinguish amino acids that contribute to function from those responsible for structure, and to focus mutations only on the former.

Computational tools are indispensable for identifying important amino acid residues in proteins, particularly for scalable and broadly applicable analyses^51, 52^. Although recent methods using NMR chemical shifts have shown promise in narrowing down potential mutation sites^53, 54^, *in silico* approaches remain central to efficiently exploring large sequence spaces. Among these, evolutionary conservation information is frequently used to estimate the functional or structural contribution of each amino acid residue^55, 56^. Highly conserved residues are likely essential for protein folding and biological activity in native contexts. However, one of the remaining challenges is separating residues critical for specific functions from those involved in structural integrity—something that conventional conservation-based methods alone cannot achieve.

In this study, we proposed a methodology in which hundreds of amino acid sequences from two structurally homologous enzyme groups are analyzed using a simple linear regression equation to extract amino acid residues where differences in substrate specificity between the enzyme groups appear. Three structurally homologous enzyme pairs (trypsin/chymotrypsin, AC/GC, and LDH/MDH) that had already been investigated in previous studies were selected, and an attempt was made to extract the amino acid residues responsible for substrate specificity from evolutionary information. By applying this method, we accurately estimated amino acid residues that are experimentally known to confer substrate specificity and identified new amino acid residues that have not yet been investigated. Furthermore, the deduced amino acid residues for the LDH/MDH pair were experimentally validated. Four LDHs from three phylogenetically distinct species were selected, and the amino acid residues identified by the ranking-based analysis were replaced with MDH-like residues to evaluate substrate specificity. The results showed that substrate specificity shifted stepwise from pyruvate to oxaloacetate as mutations accumulated. Furthermore, the expression levels of all mutants varied in the range of 40.0%–167.2%, and the introduced mutations had no significant effect on expression levels. These results suggest that the amino acid residues inferred using this method are indeed responsible for functional specificity.

We also examined the involvement of conserved amino acid residues in substrate specificity. Using the 228 LDH sequences previously employed in our EZSCAN protocol to identify substrate specificity differences between LDH and MDH, we calculated the conservation level for each residue. Highly conserved residues were distributed throughout the protein sequence (Figure 6A). Interestingly, the top five residues predicted by EZSCAN as functionally important for LDH activity had conservation ratios ranging from 0.852 to 0.988. When the conservation ratios of all residues were mapped onto the three-dimensional structure of gsLDH, we observed that the conserved residues were particularly concentrated in the structural core, the multimer interface, and the substrate-binding pocket of the protein (Figure 6B). This pattern supports earlier findings that residues in the structural core evolve more slowly than those on the surface^18^. In total, 72 residues in the gsLDH sequence had conservation ratios above 0.8, making it difficult to determine which residues were specifically responsible for substrate recognition (Figure 6C). These findings suggest that substrate-specific residues can be hidden in highly conserved regions, making them difficult to identify using conservation-based methods alone. Notably, several key residues identified using our method were not completely conserved, implying that some functionally important residues may tolerate minor evolutionary variation. This highlights the need for comparison methods that consider not only sequence conservation but also functional divergence. Our approach, which contrasts sequences based on functional differences among structurally related enzymes, offers a promising strategy for identifying such critical residues.

**Figure 6.**
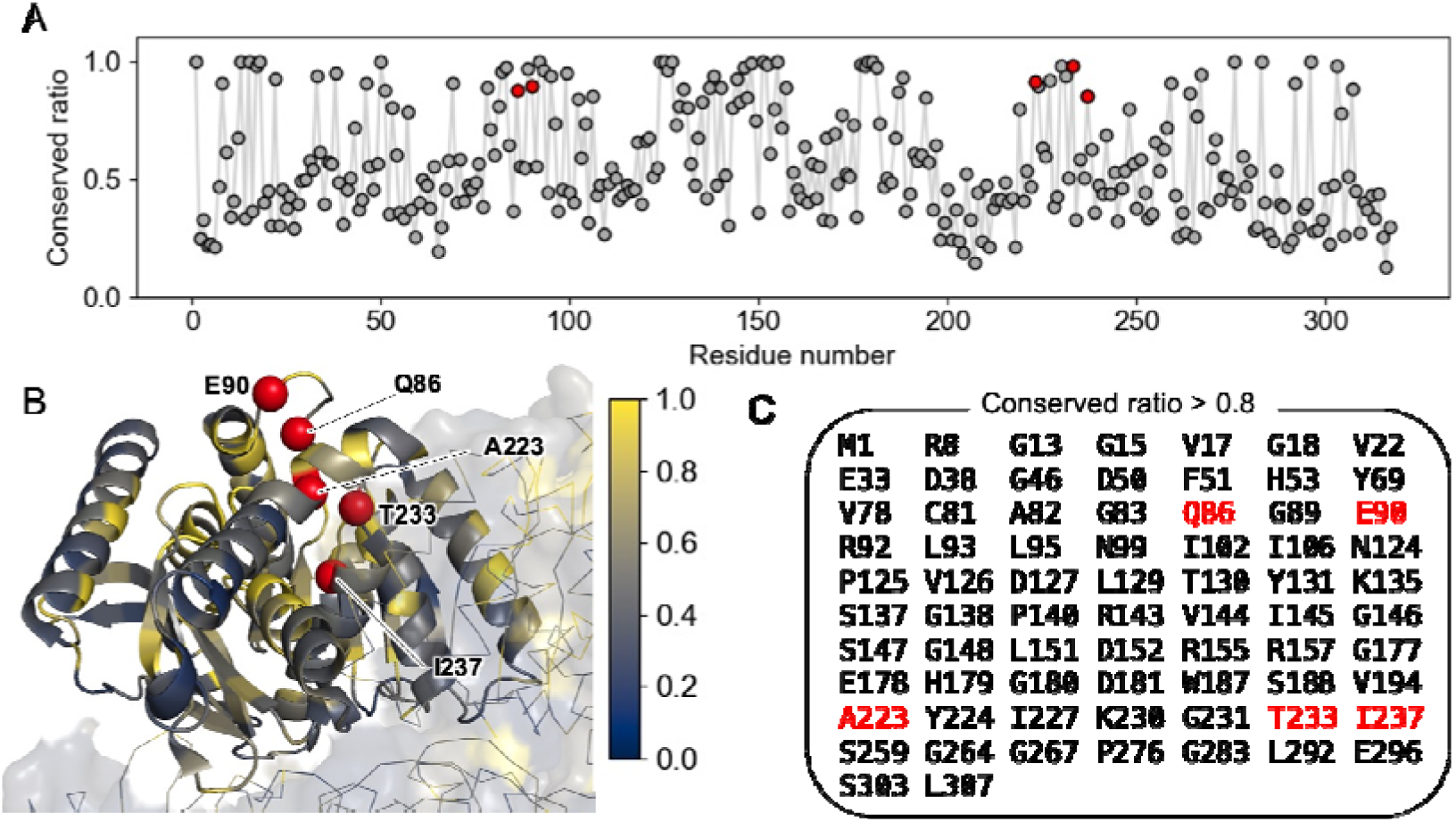
Visualization of conserved residues. (A) Conservation scores of each amino acid residue using gsLDH as the reference sequence. Scores closer to 1 indicate higher conservation. Red plots indicate the top 1–5 ranked positions predicted by the EZSCAN protocol as critical for LDH function. (B) Structural representation of gsLDH (PDB ID: 1LDB) with conservation scores mapped onto each residue. The color bar indicates the conservation ratio. (C) List of amino acid residues with conservation scores of 0.8 or higher. Residues highlighted in red represent the top 1–5 ranked positions predicted by the EZSCAN protocol as critical for LDH function.

To make this method broadly accessible, we developed a user-friendly software tool called EZSCAN. Many proteins share structural similarities despite performing different biological functions. This structural similarity can be explored using databases such as SCOP^57^, CATH^58^, and InterPro^59^, which classify proteins based on their structural and evolutionary relationships. For example, the SCOP database defines over 5,900 families of proteins that share structural motifs but often fulfill distinct biological roles. In addition to these databases, online tools such as Foldseek^60^, DALI^61^, and PDBeFold^62^ can rapidly identify proteins with similar structures. EZSCAN can be used not only to identify amino acid residues responsible for enzyme substrate specificity but also to highlight differences between other structurally similar protein pairs that differ in function. With the rapid expansion of amino acid sequence data from genome and metagenome projects, and the increasing availability of predicted protein structures driven by deep learning, the potential applications of EZSCAN are expected to grow significantly across both biology and biotechnology.

In summary, we have proposed a methodology for extracting substrate specificity-conferring residues using a linear regression-based classification program that compares groups of enzymes with homologous sequences. The estimated amino acid residues altered substrate specificity without disrupting overall protein structure. This method enables the identification and extraction of shared sequence patterns important for protein function and structure—something difficult to achieve using conservation information alone. Furthermore, because the method is highly interpretable, relying only on a linear model, it may prove useful for experimental validation of protein function and for selecting mutational targets in protein engineering.

## ASSOCIATED CONTENT

The Supporting Information is available free of charge at https://XXXXX.

Histogram of amino acid sequence length that used in the machine learning dataset (Figure S1); Aligned heatmap of specificity-conferring residues in trypsin /chymotrypsin, AC/GC, and LDH/MDH (Figure S2–S4); Structural models of pfLDH_trunc (Figure S5); SDS-PAGE image of purified LDHs and their yield (Figure S6); Substrate-dependent kinetics of LDH and MDH reactions (Figure S7); RMSD and TM-score values of pairwise structure of trypsin/chymotrypsin, AC/GC, and LDH/MDH (Table S1–S6); Kinetic parameters of LDHs (Table S7)

## AUTHOR INFORMATION

### Corresponding Authors

**Teppei Niide** − *Department of Bioinformatic Engineering, Graduate School of Information Science and Technology, The University of Osaka, 1-5 Yamadaoka, Suita, Osaka 565-0871, Japan*;

**Hiroshi Shimizu** − *Department of Bioinformatic Engineering, Graduate School of Information Science and Technology, The University of Osaka, 1-5 Yamadaoka, Suita, Osaka 565-0871, Japan*;

### Authors

**Seiya Mori** − *Department of Bioinformatic Engineering, Graduate School of Information Science and Technology, The University of Osaka, 1-5 Yamadaoka, Suita, Osaka 565-0871, Japan*

**Yoshihiro Toya** − *Department of Bioinformatic Engineering, Graduate School of Information Science and Technology, The University of Osaka, 1-5 Yamadaoka, Suita, Osaka 565-0871, Japan*

Complete contact information is available at: https://XXXXX.

### Author Contributions

T.N. and H.S. conceived and designed research, with contributions from Y.T., and S.M. conducted experiments. S.M. and T.N. analyzed and interpreted results. S.M. and T.N. wrote the manuscript with input from all authors. T.N. and H.S. supervised the study. All authors approved the content of the submitted manuscript.

### Notes

There are no conflicts of interest directly relevant to the content of this work.

## ACKNOWLEDGMENTS

We are grateful to Keiko Hiratomi and Satoko Wake for their expert technical assistance. This work was supported by Japan Society for the Promotion of Science KAKENHI (22K04841; T.N.), ACT-X (JPMJAX20BC; T.N.), PRESTO (JPMJPR24G5; T.N.), GteX (JPMJGX23B4; T.N. and Y.T.), BOOST (JPMJBS2402; S.M.) Japan Science and Technology Agency.

## Notes

### Competing Interest Statement

The authors have declared no competing interest.

