## Supporting Information for "A Method for Predicting Enzyme Substrate Specificity Residues Using Homologous Sequence Information"

This PDF file contains:

Experimental methods

Figures S1 to S7

Tables S1 to S7

Sequence information

SI References

**EXPERIMENTAL METHODS**

**Data Collection and Computational Prediction**

The amino acid sequences of trypsin, chymotrypsin, AC, GC, LDH, and MDH were collected from the UniProt^63^ and KEGG^64^ databases. The sequences were classified based on substrate selectivity and curated into non-redundant datasets for supervised machine learning. These sequences were analyzed using the scheme named the EZSCAN protocol. MSA was performed using the MAFFT^65^ and MUSCLE^66^ software to standardize sequence lengths for each enzyme pair (trypsin/chymotrypsin, AC/GC, and LDH/MDH). The aligned sequences were converted into binary representations, and each pair was assigned a label of “0” or “1” for supervised classification.

A logistic regression model was then applied, with 70% of the data used for training and 30% for testing. The partial regression coefficient (β) served as the weight parameter in the logistic function and was optimized using the steepest descent method. Specifically, the cross-entropy function was used as the loss function, and its minimization was achieved through a gradient descent algorithm. The difference between the minimum and maximum values of coefficient β was calculated and ranked in ascending order.

**Structure Similarity Analysis**

The crystal structures of trypsin, chymotrypsin, AC, GC, LDH, and MDH were obtained from the Protein Data Bank. The PDB IDs for these crystal structures are listed in the headers of Supplementary Tables S1–S7. Structural alignment for each enzyme pair was conducted using TM-align^67^. The aligned structures, along with the corresponding RMSD and TM-score, were outputs generated by TM-align.

**Phylogenetic Analysis**

The amino acid sequences of LDH and MDH used in the EZSCAN protocol were employed for phylogenetic analysis. MSA was performed using MAFFT, ^65^ and poorly aligned regions were removed. The phylogenetic tree was constructed using the maximum likelihood method in IQ-Tree 2^68^. The Bayesian information criterion with ModelFinder ^69^ was used for selecting the substitution, with the LG+R9 model being selected. The reliability of the estimated clade was evaluated using the bootstrap method with UFBoot2 ^70^ and 1500 bootstrap iterations.

**Plasmid Construction**

Wild-type lcLDH, gsLDH, pfLDH, and their corresponding mutants were synthesized using the GeneArt gene synthesis service (Thermo Fisher Scientific, Waltham, MA, USA). Each DNA fragment was amplified via PCR using KOD polymerase (Toyobo, Osaka, Japan) and subsequently purified. The genes and the pET28a vector were digested with NdeI and XhoI restriction enzymes (New England Biolabs, Ipswich, MA, USA), followed by ligation into the linearized pET28a vector using T4 DNA ligase (Toyobo). The pfLDH_trunc expression plasmids were constructed using inverse PCR and a self-ligation method with the pfLDH expression plasmids as templates. All strains used in this study were selected on LB agar plates supplemented with 30 µg mL⁻¹ kanamycin. The integrity of the gene sequences was verified by Sanger sequencing.

**Protein Expression and Purification**

*E. coli* strain BL21(DE3) was transformed with the constructed plasmids and plated on LB agar containing 30 µg mL⁻¹ kanamycin. A single colony was randomly selected and grown in LB liquid medium with 30 µg mL⁻¹ kanamycin at 37°C. Following overnight incubation, the culture was transferred into baffled Erlenmeyer flasks containing 2× YT medium with 30 µg mL⁻¹ kanamycin and grown at 37°C with shaking. When the optical density at 600 nm reached 0.4–0.6, IPTG was added to a final concentration of 0.5 mM to induce protein expression, and the culture was incubated overnight at 18°C with shaking.

Cells were harvested by centrifugation and lysed using BugBuster Master Mix (Merck Millipore, Burlington, MA, USA). After gently stirring at room temperature for 20 min, the lysate was centrifuged, and the supernatant was collected. The supernatants were filtered through 0.22 µm pore size membranes (Merck, Kenilworth, NJ, USA) and applied to Ni-NTA resin-packed columns for purification. The resin was washed with Buffer A (50 mM Tris-HCl, 200 mM NaCl, and 50 mM imidazole) and eluted with Buffer B (50 mM Tris-HCl, 200 mM NaCl, and 300 mM imidazole). The eluted proteins were subjected to buffer exchange into Buffer C (50 mM Tris-HCl and 200 mM NaCl) using a PD-10 column (Cytiva, Marlborough, MA, USA). Protein concentrations were determined by measuring absorbance at 280 nm.

**Kinetic Assay**

Enzyme activity was assessed by monitoring the dynamic consumption of NADH during the reductive conversion of pyruvate to lactate and oxaloacetate to malate, measured at 340 nm using a Synergy HTX multi-mode reader (Agilent Technologies Inc., Santa Clara, CA, USA). Each reaction was performed in a 200 µL volume, and measurements were taken for 10 min, with the enzyme solution added last to initiate the reaction.

The reaction mixture contained 0.3 mM NADH, 50 mM Tris-HCl (pH 7.4) for wild-type gsLDH, pfLDH, pfLDH_trunc, and their variants, or 50 mM acetate buffer (pH 4.8) for wild-type lcLDH and its variants. The mixture also included 3 mM fructose-1,6-bisphosphate as an activator^71^ and 0.01–35 mM sodium pyruvate or 0.01–4 mM oxaloacetate as the substrate. The rate of substrate conversion was calculated based on changes in NADH absorbance. Kinetic parameters were derived from the average values of three independent replicates. The reaction rate was determined from the slope of the linear portion of the absorbance curve and fitted to the Michaelis–Menten model using Python.

**SUPPLEMENTARY FIGURES**

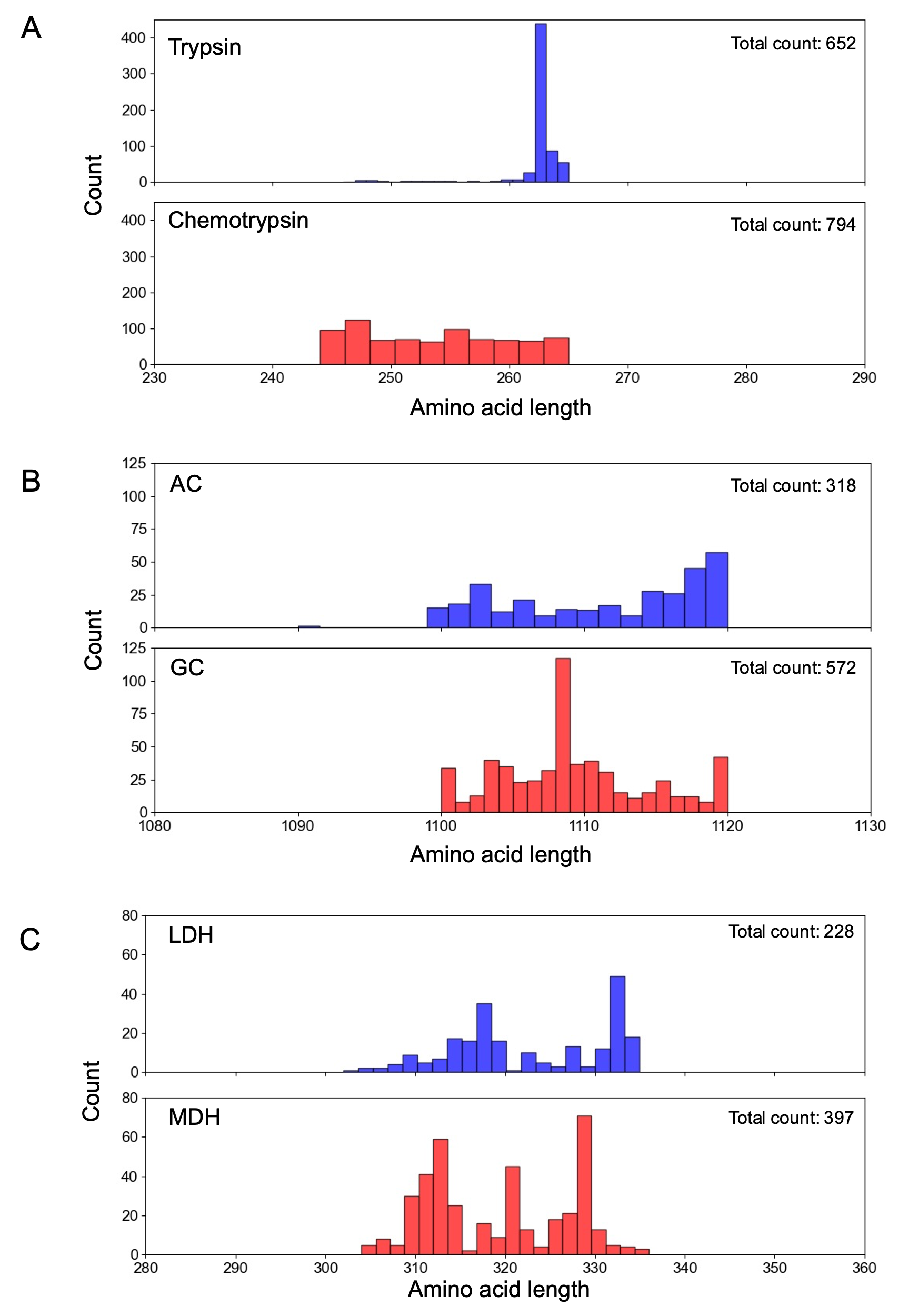

Figure S1. Histogram of amino acid sequence length that used in the machine learning dataset. (A) Trypsin (blue) and chymotrypsin (red). (B) AC (blue) and GC (red). (C) LDH (blue) and MDH (red). It contains a total of 952 amino acid sequences randomly obtained from the KEGG database

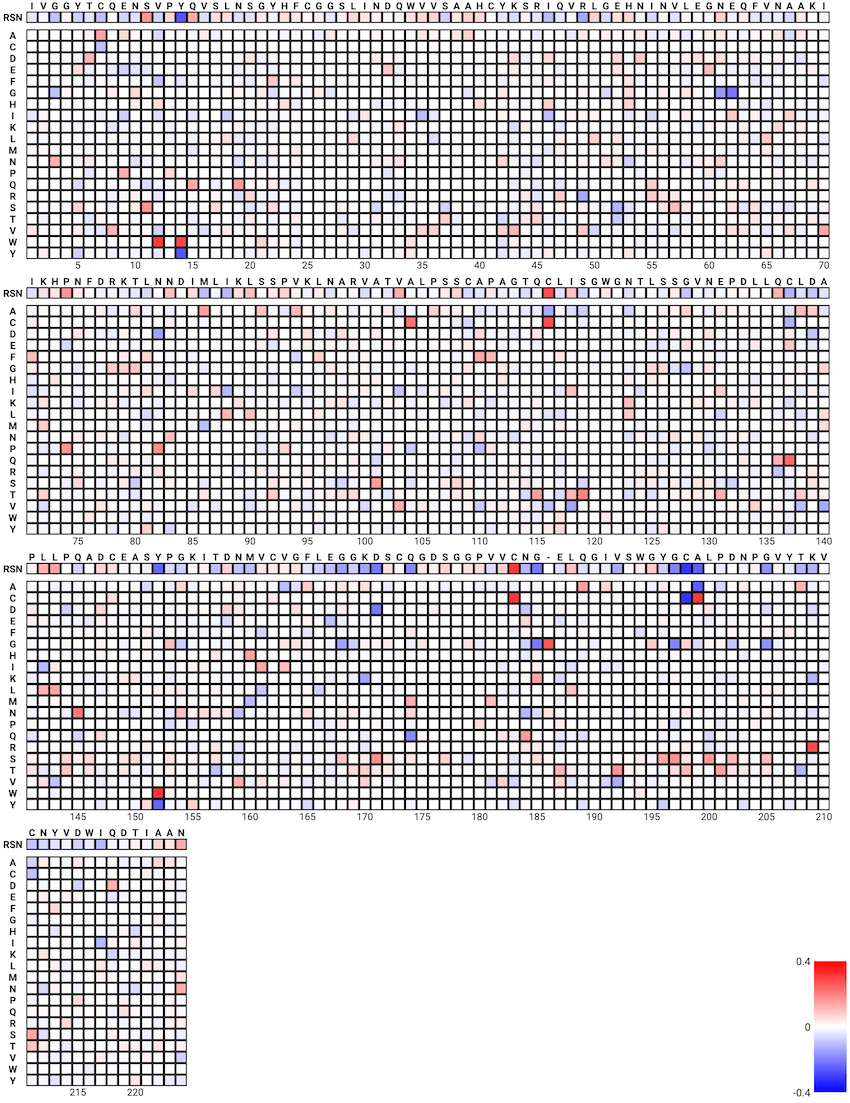

**0.4**

**Trypsin**

**Chymotrypsin**

**−0.4**

**0**

Figure S2. Aligned heatmap showing the properties of amino acid residues involved in substrate specificity of trypsin and chymotrypsin. Blue and red indicate residues highly associated with trypsin and chymotrypsin, respectively. RSN refers to *Rattus norvegicus* trypsin, with each residue colored according to its contribution to substrate specificity based on the same heatmap scale. Columns enclosed by hyphens in the *R. norvegicus* trypsin sequence indicate the top 30 ranked residues most strongly associated with substrate specificity according to the model.

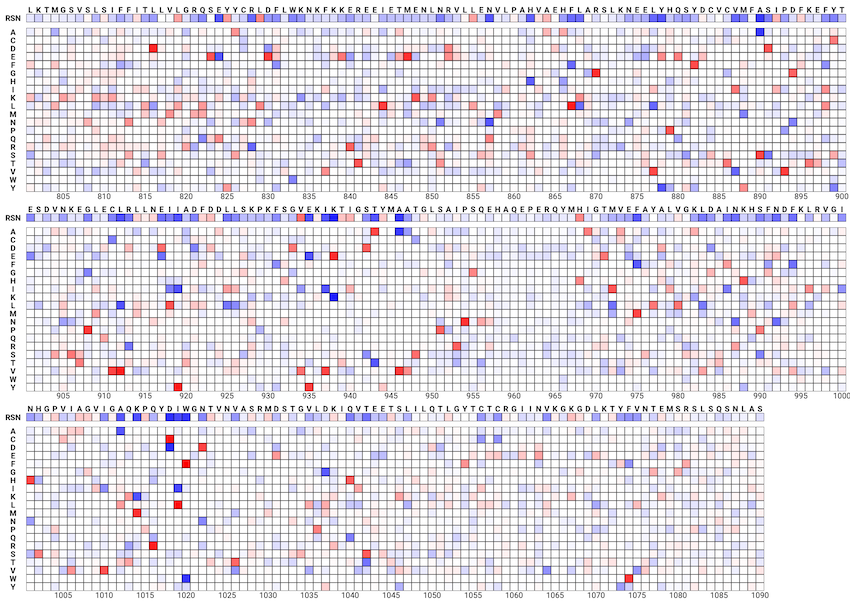

**0.05**

**AC**

**GC**

**−0.05**

**0**

Figure S3. Aligned heatmap illustrating the residue features associated with the functional specificity of AC and GC, focusing on positions beyond residue 800, which include the catalytic C2 domain. Blue and red indicate residues strongly associated with AC and GC, respectively. The panel labeled RSN corresponds to *R. norvegicus* AC, with a similar heatmap showing the contribution of each residue to substrate specificity.

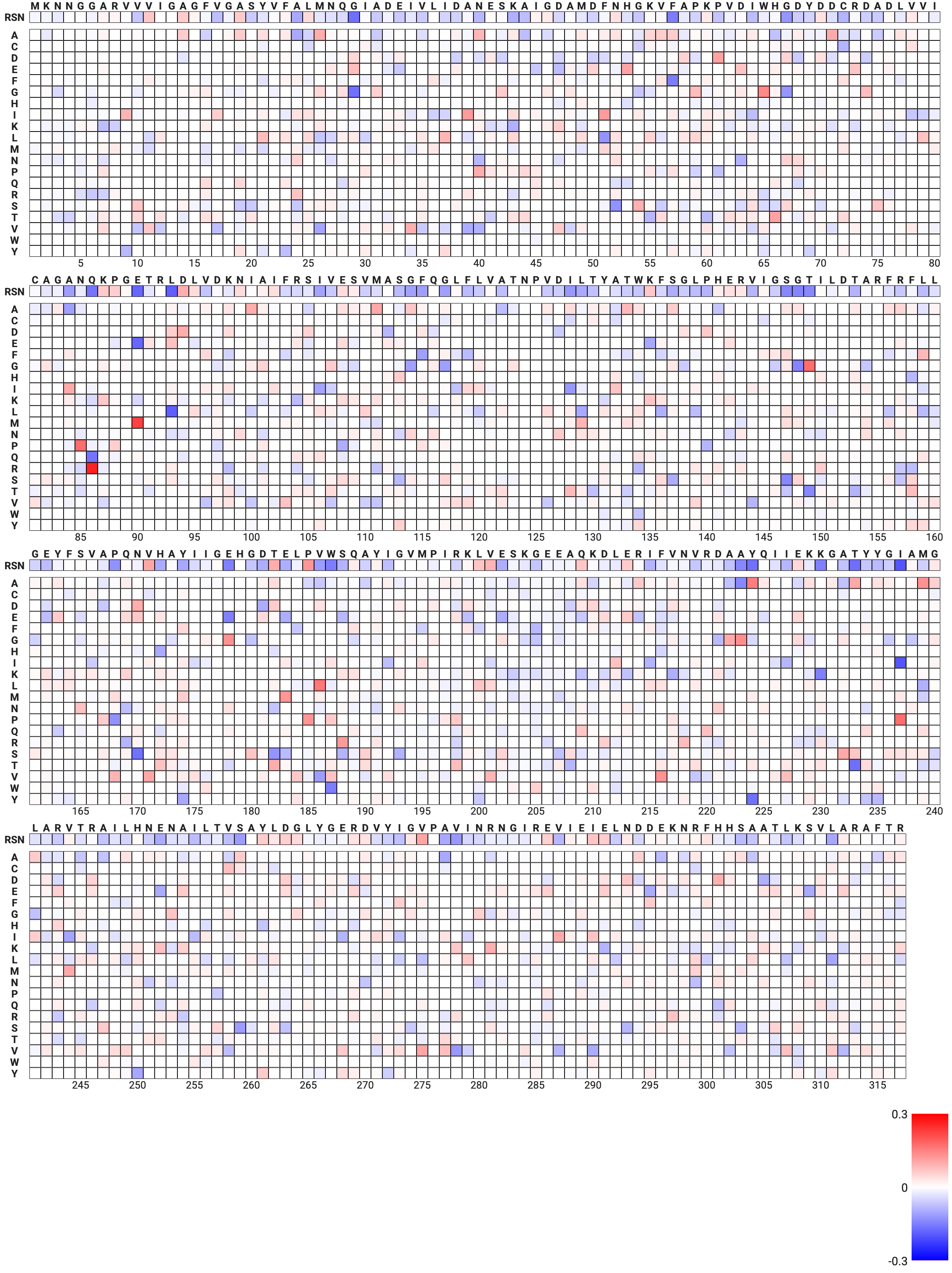

**0.3**

**LDH**

**MDH**

**−0.3**

**0**

Figure S4. Aligned heatmap illustrating the residue features associated with the functional specificity of LDH and MDH. Blue and red indicate residues strongly associated with LDH and MDH, respectively. The panel labeled RSN corresponds to *Geobacillus stearothermophilus* LDH, with a similar heatmap showing the contribution of each residue to substrate specificity.

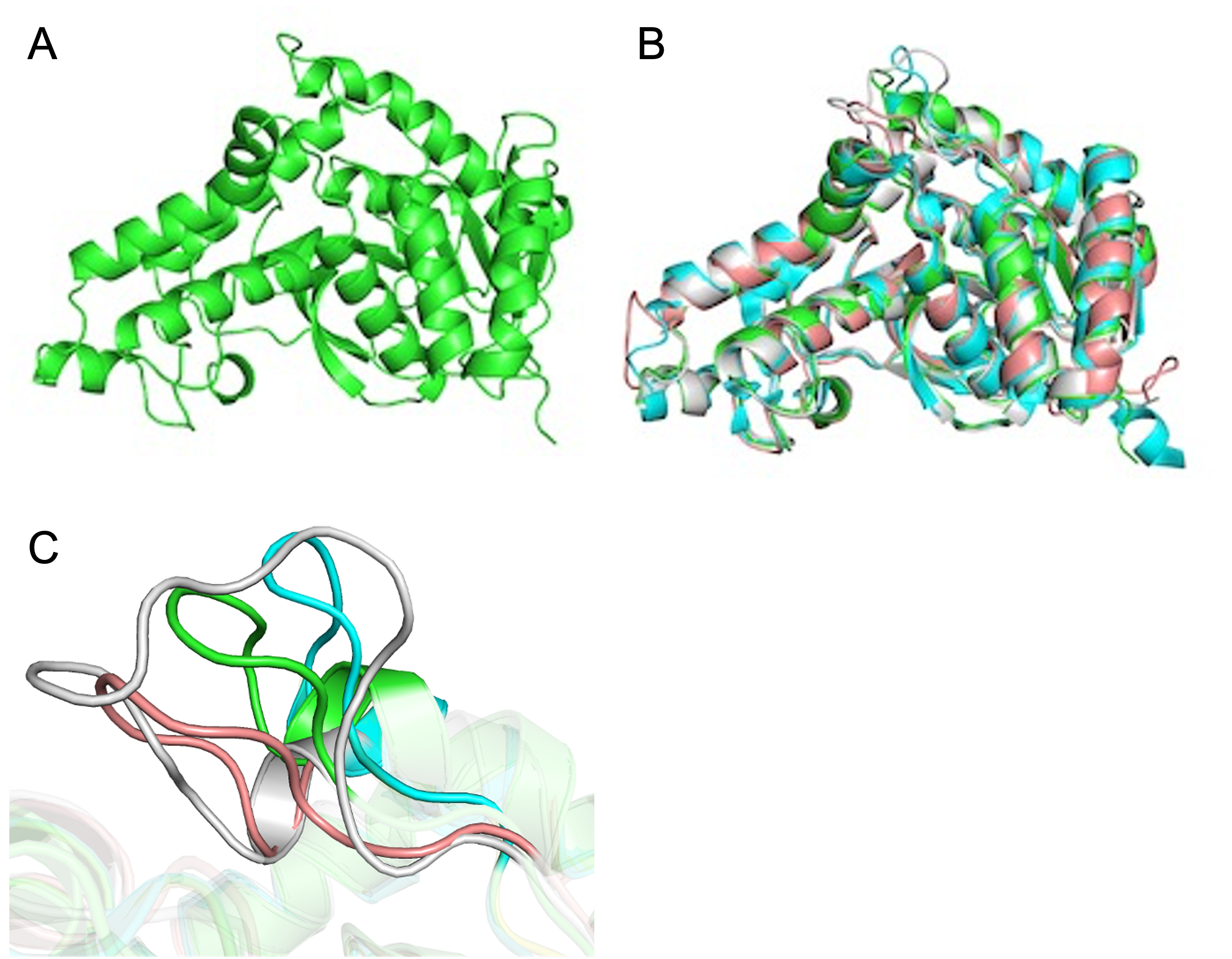

Figure S5. (A) Predicted structure of pfLDH_trunc generated by AlphaFold2. (B) Superimposed image of the predicted pfLDH_trunc structure (green) with the crystal structures of pfLDH (gray), gsLDH (salmon), and lcLDH (cyan). (C) Close-up view of the superimposed structures highlighting the region F82–D97 residues of pfLDH, which are involved in substrate specificity.

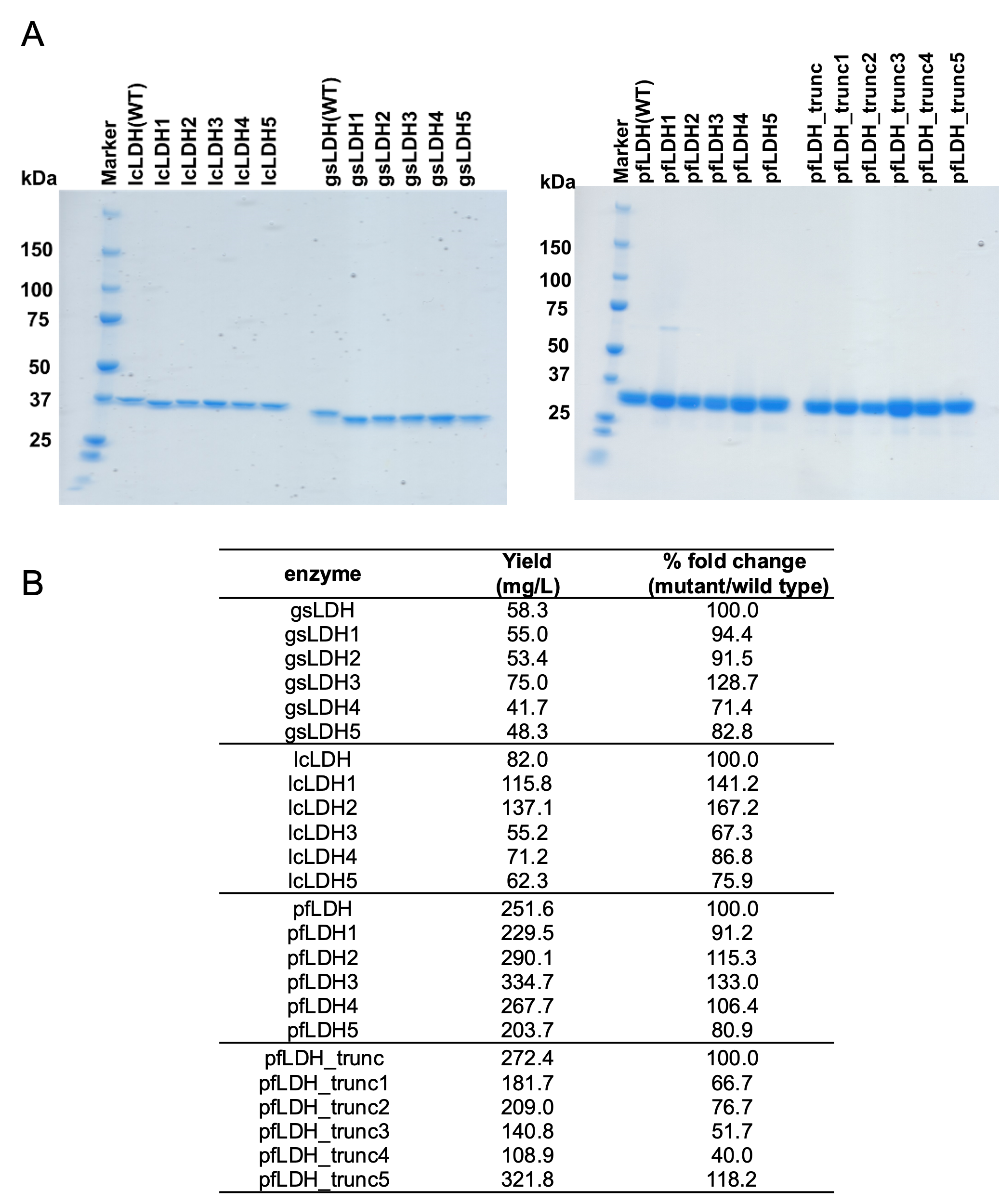

Figure S6. (A) SDS-PAGE analysis of purified lcLDH, gsLDH, pfLDH, pfLDH_trunc, and their respective mutants obtained by metal affinity chromatography. (B) Yields of each purified protein.

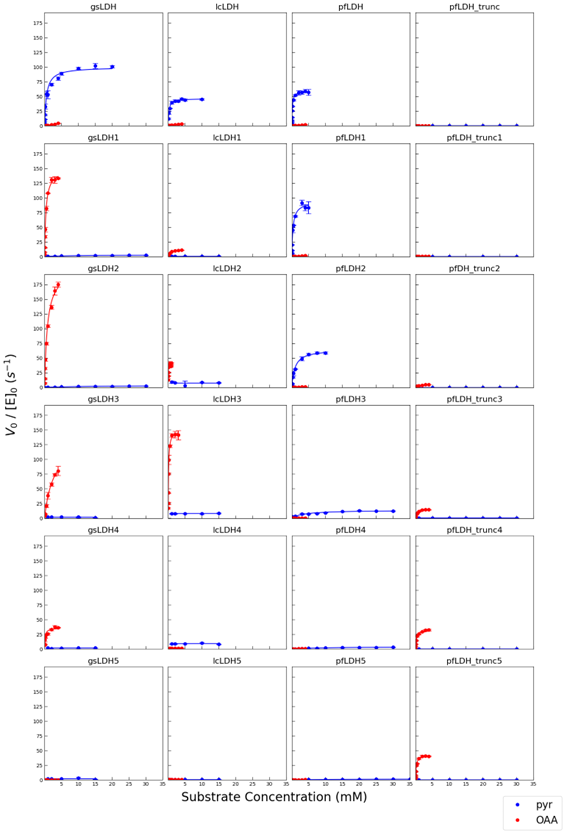

Figure S7. Substrate concentration dependence of the initial reaction velocity. Pyruvate (blue dots and line), the substrate for LDH, and oxaloacetate (red dots and line), the substrate for MDH, were used. Nonlinear fitting to the Michaelis–Menten model was performed to calculate kinetic parameters, which are summarized in Table S7.

**SUPPLEMENTARY TABLES**

Table S1. RMSD values of each pairwise structure in the list of trypsin/chymotrypsin.

| PDB ID | 1ACB | 1ANE | 1EQ9 | 1KDQ | 1OS8 | 6T5W |
| --- | --- | --- | --- | --- | --- | --- |
| 1ACB | 0 |  |  |  |  |  |
| 1ANE | 1.21 | 0 |  |  |  |  |
| 1EQ9 | 1.5 | 1.24 | 0 |  |  |  |
| 1KDQ | 1.34 | 1.59 | 1.83 | 0 |  |  |
| 1OS8 | 1.63 | 1.44 | 1.85 | 2.06 | 0 |  |
| 6T5W | 1.25 | 0.41 | 1.26 | 1.59 | 1.46 | 0 |

Table S2. TM-score values of each pairwise structure in the list of trypsin/chymotrypsin.

| PDB ID | 1ACB | 1ANE | 1EQ9 | 1KDQ | 1OS8 | 6T5W |
| --- | --- | --- | --- | --- | --- | --- |
| 1ACB | 1 |  |  |  |  |  |
| 1ANE | 0.946 | 1 |  |  |  |  |
| 1EQ9 | 0.924 | 0.931 | 1 |  |  |  |
| 1KDQ | 0.957 | 0.891 | 0.872 | 1 |  |  |
| 1OS8 | 0.893 | 0.897 | 0.882 | 0.471 | 1 |  |
| 6T5W | 0.944 | 0.995 | 0.925 | 0.532 | 0.896 | 1 |

Table S3. RMSD values of each pairwise structure in the list of AC/GC.

| PDB ID | 1AB8 | 2WZ1 | 3ET6 | 6PAS | 6R3Q | 7YZI |
| --- | --- | --- | --- | --- | --- | --- |
| 1AB8 | 0 |  |  |  |  |  |
| 2WZ1 | 2.29 | 0 |  |  |  |  |
| 3ET6 | 2.28 | 1.60 | 0 |  |  |  |
| 6PAS | 2.82 | 2.64 | 2.69 | 0 |  |  |
| 6R3Q | 1.64 | 2.11 | 2.04 | 5.76 | 0 |  |
| 7YZI | 1.92 | 1.74 | 1.96 | 3.83 | 3.82 | 0 |

Table S4. TM-score values of each pairwise structure in the list of AC/GC.

| PDB ID | 1AB8 | 2WZ1 | 3ET6 | 6PAS | 6R3Q | 7YZI |
| --- | --- | --- | --- | --- | --- | --- |
| 1AB8 | 1 |  |  |  |  |  |
| 2WZ1 | 0.835 | 1 |  |  |  |  |
| 3ET6 | 0.805 | 0.855 | 1 |  |  |  |
| 6PAS | 0.750 | 0.770 | 0.801 | 1 |  |  |
| 6R3Q | 0.911 | 0.840 | 0.886 | 0.393 | 1 |  |
| 7YZI | 0.856 | 0.804 | 0.838 | 0.373 | 0.305 | 1 |

Table S5. RMSD values of each pairwise structure in the list of LDH/MDH.

| PDB ID | 1B8P | 1HLP | 1LDG | 1LDN | 2PWZ | 3VPH | 4AJ2 | 4CL3 | 5UJK | 6J9T |
| --- | --- | --- | --- | --- | --- | --- | --- | --- | --- | --- |
| 1B8P | 0 |  |  |  |  |  |  |  |  |  |
| 1HLP | 2.44 | 0 |  |  |  |  |  |  |  |  |
| 1LDG | 2.50 | 2.03 | 0 |  |  |  |  |  |  |  |
| 1LDN | 2.69 | 1.99 | 1.57 | 0 |  |  |  |  |  |  |
| 2PWZ | 2.45 | 2.26 | 2.32 | 2.45 | 0 |  |  |  |  |  |
| 3VPH | 2.51 | 1.91 | 1.88 | 2.13 | 2.20 | 0 |  |  |  |  |
| 4AJ2 | 2.75 | 2.03 | 1.89 | 1.97 | 2.47 | 2.05 | 0 |  |  |  |
| 4CL3 | 2.32 | 1.82 | 1.56 | 1.79 | 2.10 | 1.59 | 2.09 | 0 |  |  |
| 5UJK | 2.26 | 1.84 | 1.50 | 1.99 | 2.09 | 1.71 | 2.28 | 1.03 | 0 |  |
| 6J9T | 2.36 | 1.77 | 1.67 | 1.77 | 2.35 | 1.45 | 1.88 | 1.40 | 1.45 | 0 |

Table S6. TM-score values of each pairwise structure in the list of LDH/MDH.

| PDB ID | 1B8P | 1HLP | 1LDG | 1LDN | 2PWZ | 3VPH | 4AJ2 | 4CL3 | 5UJK | 6J9T |
| --- | --- | --- | --- | --- | --- | --- | --- | --- | --- | --- |
| 1B8P | 1 |  |  |  |  |  |  |  |  |  |
| 1HLP | 0.820 | 1 |  |  |  |  |  |  |  |  |
| 1LDG | 0.830 | 0.911 | 1 |  |  |  |  |  |  |  |
| 1LDN | 0.831 | 0.922 | 0.937 | 1 |  |  |  |  |  |  |
| 2PWZ | 0.820 | 0.863 | 0.828 | 0.821 | 1 |  |  |  |  |  |
| 3VPH | 0.821 | 0.923 | 0.910 | 0.922 | 0.840 | 1 |  |  |  |  |
| 4AJ2 | 0.833 | 0.919 | 0.914 | 0.943 | 0.839 | 0.922 | 1 |  |  |  |
| 4CL3 | 0.840 | 0.924 | 0.925 | 0.913 | 0.860 | 0.928 | 0.874 | 1 |  |  |
| 5UJK | 0.849 | 0.923 | 0.945 | 0.924 | 0.861 | 0.936 | 0.873 | 0.978 | 1 |  |
| 6J9T | 0.828 | 0.917 | 0.919 | 0.938 | 0.820 | 0.938 | 0.873 | 0.924 | 0.920 | 1 |

**Table S7** Kinetic parameters of wild-type and mutant LDHs. The number associated with each mutant indicates the total number of amino acid substitutions. LDH activity was measured using pyruvate as the substrate, while MDH-like activity was evaluated using oxaloacetate, enabling comparison of catalytic efficiency across variants.

|  | K_M_ (mM) | | k_cat_ (s^-1^) | | k_cat_/K_M_ (mM^-1^s^-1^) | |
| --- | --- | --- | --- | --- | --- | --- |
|  | pyruvate | oxaloacetate | pyruvate | oxaloacetate | pyruvate | oxaloacetate |
| gsLDH | 0.57 ± 0.01 | n.d | 99.8 ± 0.0 | n.d | 175.1 ± 3.1 | n.d |
| gsLDH1 | 7.63 ± 0.66 | 0.41 ± 0.00 | 2.9 ± 0.0 | 150.1 ± 0.0 | 0.4 ± 0.0 | 366.1± 0.0 |
| gsLDH2 | 16.26 ± 5.00 | 0.85 ± 0.02 | 3.6 ± 0.0 | 205.5 ± 0.0 | 0.2 ± 0.1 | 241.8 ± 5.7 |
| gsLDH3 | n.d | 2.69 ± 0.07 | n.d | 136.8 ± 0.0 | n.d | 50.9 ± 1.4 |
| gsLDH4 | n.d | 0.16 ± 0.00 | n.d | 35.8 ± 0.0 | n.d | 223.8 ± 0.0 |
| gsLDH5 | n.d | n.d | n.d | n.d | n.d | n.d |
| lcLDH | 0.21 ± 0.00 | 3.27 ± 2.23 | 46.3 ± 0.0 | 5.6 ± 0.0 | 220.5 ± 0.0 | 1.7 ± 1.2 |
| lcLDH1 | n.d. | 0.48 ± 0.01 | n.d. | 12.2 ± 0.0 | n.d. | 25.4± 0.5 |
| lcLDH2 | n.d. | 0.02 ± 0.00 | n.d. | 40.5 ± 0.0 | n.d. | 2025.0± 0.0 |
| lcLDH3 | n.d. | 0.11 ± 0.00 | n.d. | 150.3 ± 0.0 | n.d. | 1336.4± 0.0 |
| lcLDH4 | n.d. | 0.09 ± 0.00 | n.d. | 1.6 ± 0.0 | n.d. | 17.8± 0.0 |
| lcLDH5 | n.d. | n.d. | n.d. | n.d. | n.d. | n.d. |
| pfLDH | 0.15 ± 0.00 | n.d. | 59.8 ± 0.0 | n.d. | 398.7 ± 0.0 | n.d. |
| pfLDH1 | 0.35 ± 0.00 | n.d. | 93.5 ± 0.0 | n.d. | 267.1 ± 0.0 | n.d. |
| pfLDH2 | 0.91 ± 0.00 | n.d. | 64.7 ± 0.0 | n.d. | 71.1 ± 0.0 | n.d. |
| pfLDH3 | 3.52 ± 0.99 | n.d. | 13.6 ± 0.0 | n.d. | 3.9 ± 1.1 | n.d. |
| pfLDH4 | 10.35 ± 0.72 | 8.35 ± 1.87 | 4.0 ± 0.0 | 2.5 ± 0.0 | 0.4 ± 0.0 | 0.3 |
| pfLDH5 | 20.37 ± 3.31 | n.d. | 1.7 ± 0.0 | n.d. | 0.1 ± 0.0 | n.d. |
| pfLDH_trunc | n.d. | n.d. | n.d. | n.d. | n.d. | n.d. |
| pfLDH_trunc1 | n.d. | 0.61 ± 0.07 | n.d. | 1.0 ± 0.0 | n.d. | 1.6 ± 0.2 |
| pfLDH_trunc2 | n.d. | 1.10 ± 0.02 | n.d. | 6.1 ± 0.0 | n.d. | 5.5 ± 0.1 |
| pfLDH_trunc3 | n.d. | 0.51 ± 0.00 | n.d. | 16.8 ± 0.0 | n.d. | 32.9 ± 0.0 |
| pfLDH_trunc4 | n.d. | 0.30 ± 0.00 | n.d. | 34.3 ± 0.0 | n.d. | 114.3 ± 0.0 |
| pfLDH_trunc5 | n.d. | 0.23 ± 0.00 | n.d. | 43.3 ± 0.0 | n.d. | 188.3 ± 0.0 |
| *"n.d." not determined due to the inability to fit | | | | | | |

**SEQUENCE INFORMATION**

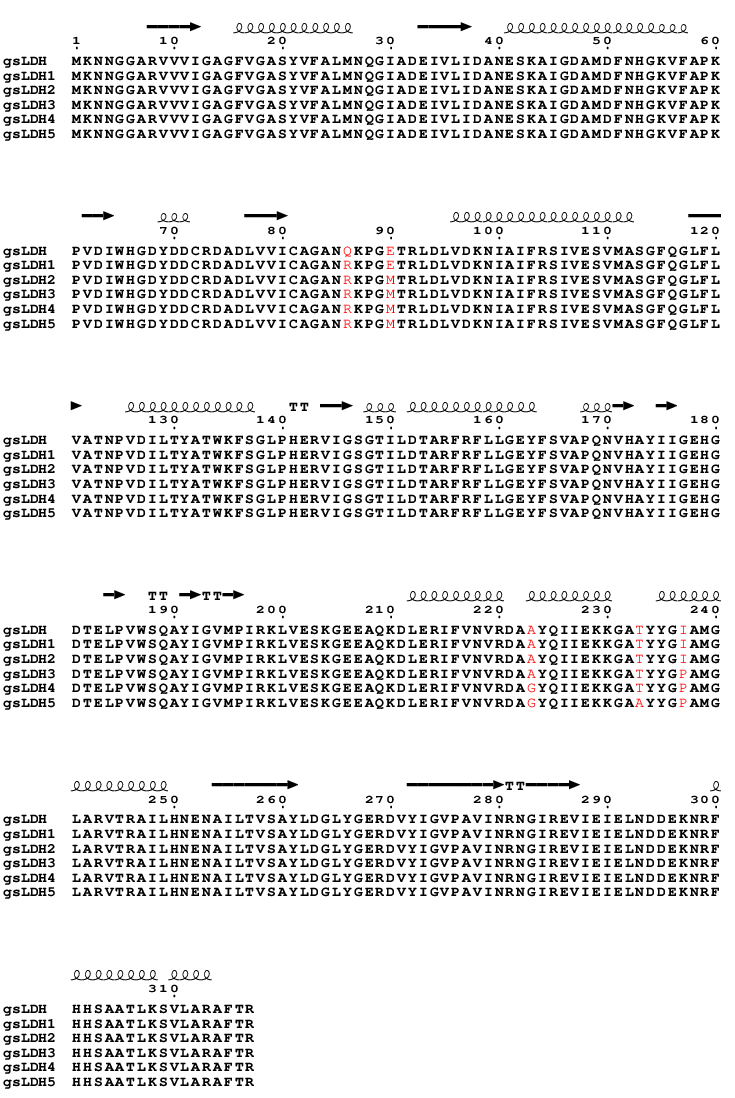

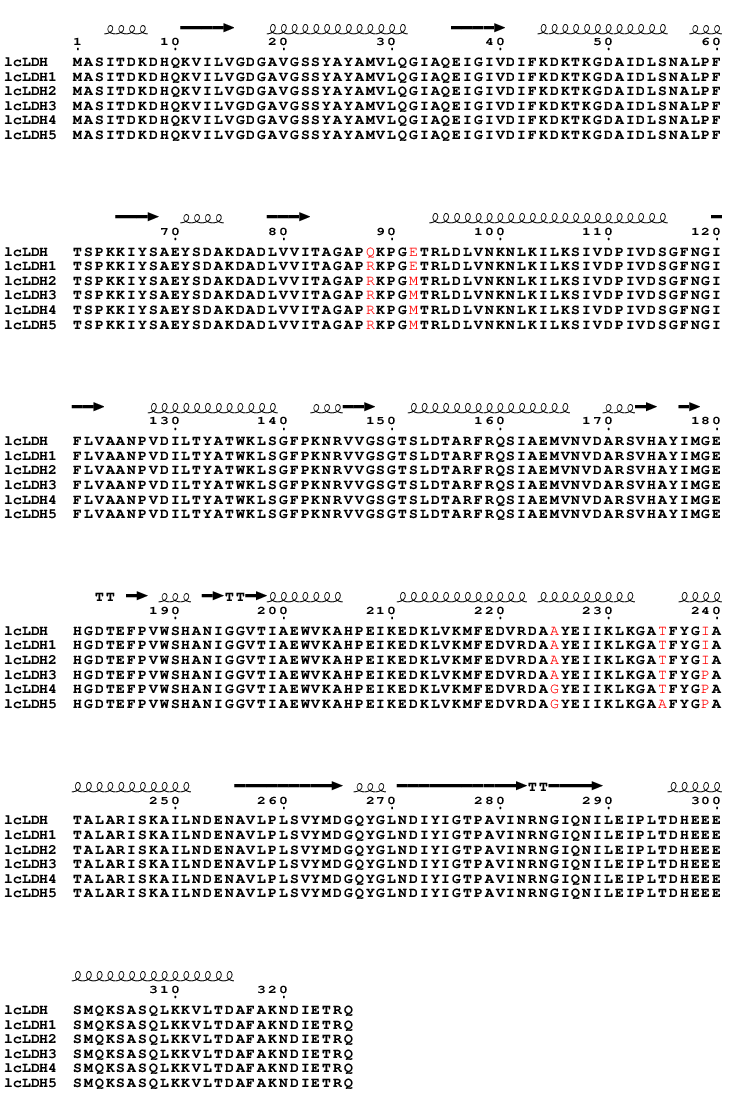

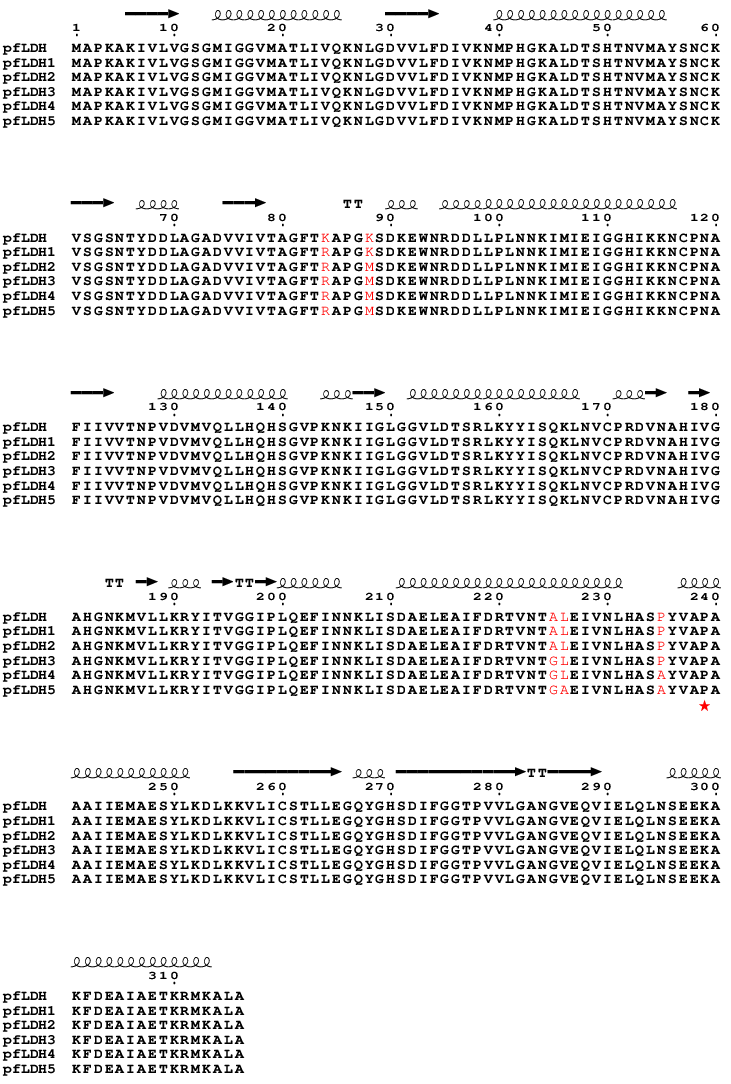

Red star mark: Although Pro239 ranked third, it already exhibited MDH-like.

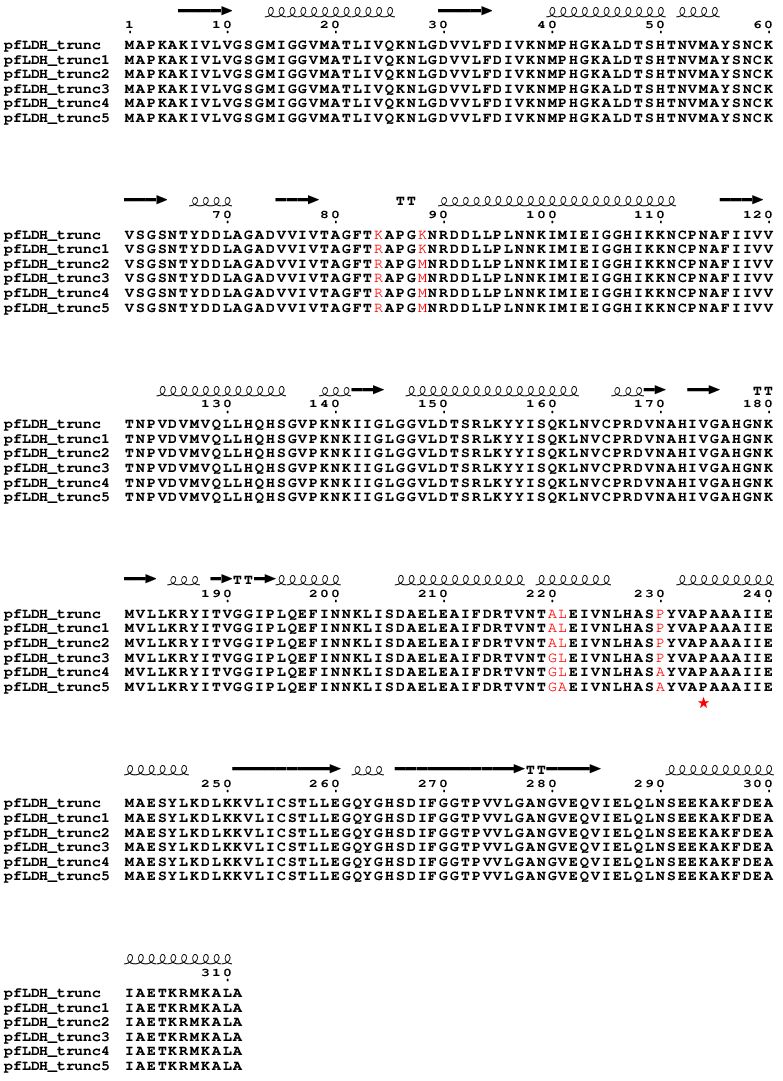

Red star mark: Although Pro234 ranked third, it already exhibited MDH-like.
